# Rapid and Reproducible Multimodal Biological Foundation Model Development with AIDO.ModelGenerator

**DOI:** 10.1101/2025.06.30.662437

**Authors:** Caleb N. Ellington, Dian Li, Shuxian Zou, Elijah Cole, Ning Sun, Sohan Addagudi, Le Song, Eric P. Xing

## Abstract

Foundation models (FMs) for DNA, RNA, proteins, cells, and tissues have begun to close long-standing performance gaps in biological prediction tasks, yet each modality is usually studied in isolation. Bridging them requires software that can ingest heterogeneous data, apply large pre-trained backbones from various sources, and perform multimodal benchmarking studies at scale. We present AIDO.ModelGenerator, an open-source toolkit that turns these needs into declarative experiment recipes through a structured experimental framework. AIDO.ModelGenerator provides (i) 300+ datasets covering DNA, RNA, protein, cell, spatial, and multimodal data types; (ii) 30+ pretrained FMs ranging from 3M to 16B parameters; (iii) 10+ plug-and-play use-cases covering inference, adaptation, prediction, generation, and zero-shot evaluation; and (iv) YAML-driven experiment recipes that enable exact reproducibility. On a sequence-to-expression prediction task, AIDO.ModelGenerator systematically builds and tests unimodal and multimodal models, achieving a new SOTA by combining DNA and RNA FMs that outperforms unimodal baselines by over 10%. In a Crohn’s disease case-study, the framework’s simulated knockout protocol ranks the clinically implicated target SOX4 6,000 positions higher than differential-expression baselines, illustrating its utility for therapeutic target discovery. We release code, tutorials, checkpoints, datasets, and API reference to accelerate multimodal FM research in the life sciences^1^.

## 1. Introduction

Self-supervised pre-training on vast collections of biological data has produced FMs capable of capturing the syntax and semantics of DNA, RNA, proteins, small molecules, single-cell states, and tissue images. In many domains, these models routinely surpass hand-engineered architectures while requiring far fewer labeled examples to be adapted for down-stream tasks. Despite this promise, day-to-day adoption in academic and industrial labs remains slow, owing to three persistent challenges.

### Heterogeneous data and fragile tooling

Biomedical research pipelines depend on dozens of data formats and modalities, each handled by a different library and tokenizer. Seemingly innocuous changes in canonicalization (e.g. ambiguous nucleotides, degenerate amino acids) or tool versioning can eliminate expected gains, and reproducing an environment months later is often impossible once Python wheels, CUDA toolkits, and random seeds have drifted.

### Computational scale and engineering burden

Modern pretrained biomolecular FMs range from 0.1B to 100B parameters. Finetuning these models requires custom architectural adaptaions as well as systems optimizations such as mixed precision, distributed optimizers, and non-trivial learning-rate schedules–skills that most wet-lab groups do not possess. While commercial cloud instances democratize access to GPUs, the software scaffolding for fault-tolerant training, checkpoint sharding, and experiment tracking remains a barrier.

### Multimodal fusion

Biological questions rarely respect modality boundaries: understanding tissue-specific expression, mapping disease pathways, or ranking therapeutic targets requires complex cross-modality workflows that often confound iterative engineering and case-control study design. Additionally, FMs each contain many idiosyncrasies and design choices, such as encoders, tokenizers, and loss functions. Even simple approaches to fusing models can require substantial engineering overhead. This difficulty compounds with the number of FMs being used, and a common experimental framework is currently missing.

Several open-source efforts have provided DL-enabled tool-ing for specific modalities, focusing on inference and fine-tuning of fixed domain-specific models and architectures (Lal et al., 2024; Ramsundar et al., 2019; Chen et al., 2019; noa; Cardoso et al., 2022). BioNeMo (John et al., 2024) extends this, freezing popular pretrained models from multiple modalities to provide efficient inference and finetuning capabilities. BioNeMo models are optimized for efficiency, prohibiting substantial architectural changes or composing new models from new pretrained components.

Experimenting or merging models in a way that changes gradient flow currently requires hands-on use of low-level scaffolding, such as Hugging Face (Lhoest et al., 2021; Wolf et al., 2020) or PyTorch (Paszke et al., 2019), which is time-consuming and must be hand-tailored to each model. Hugging Face and bio-related wrappers such as MultiMolecule (Chen & Zhu, 2024) provide some capabilities for adapting individual models with simple heads, but only for basic supervised tasks and only on text data.

While these ecosystems markedly lower the barrier for single-modal or task-specific work, none yet offer a workflow that can assist with building and benchmarking foundation models across multiple data modalities.

### 1.1. Our Contribution

We present **AIDO.ModelGenerator**, an open-source toolkit built on PyTorch Lightning and Hugging Face for multi-modal FM usage, benchmarking, and development. The framework provides

1. standard interfaces for integrating FMs, applications, and datasets, such that these components can be (i) rapidly swapped and combined and (ii) developed independently of one another;
2. declarative YAML “recipes” that compile into fully version-pinned, container-ready experiments;
3. a curated suite of 300+ benchmarks and 30+ back-bones, as well as no-code entrypoints for working with external models and datasets.
4. automatic scripts for leaderboard creation and hyper-parameter tuning; and
5. first-class distributed training, parameter-efficient fine-tuning, and automatic experiment tracking.

## 2. Methods

AIDO.ModelGenerator achieves interoperability of datasets, backbones, tasks, and adapters through interfaces which wrap these components in standard APIs (Fig. 1). These components are automatically compiled together by passing a declarative YAML configuration to the `mgen` command.

**Figure 1.**
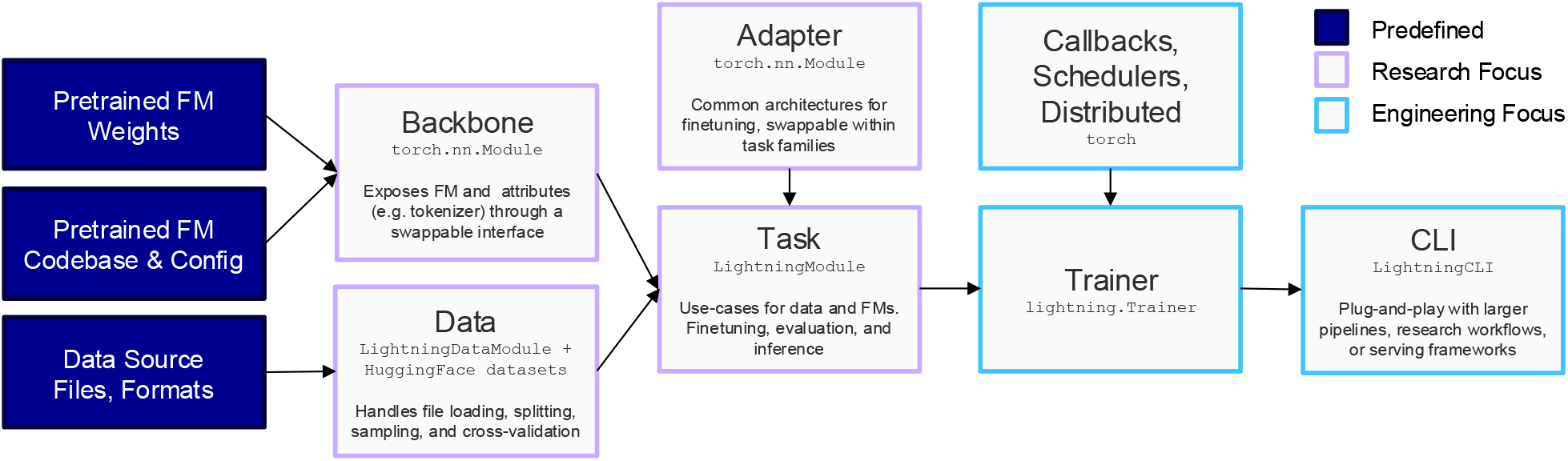
AIDO.ModelGenerator architecture.

**Figure 2.**
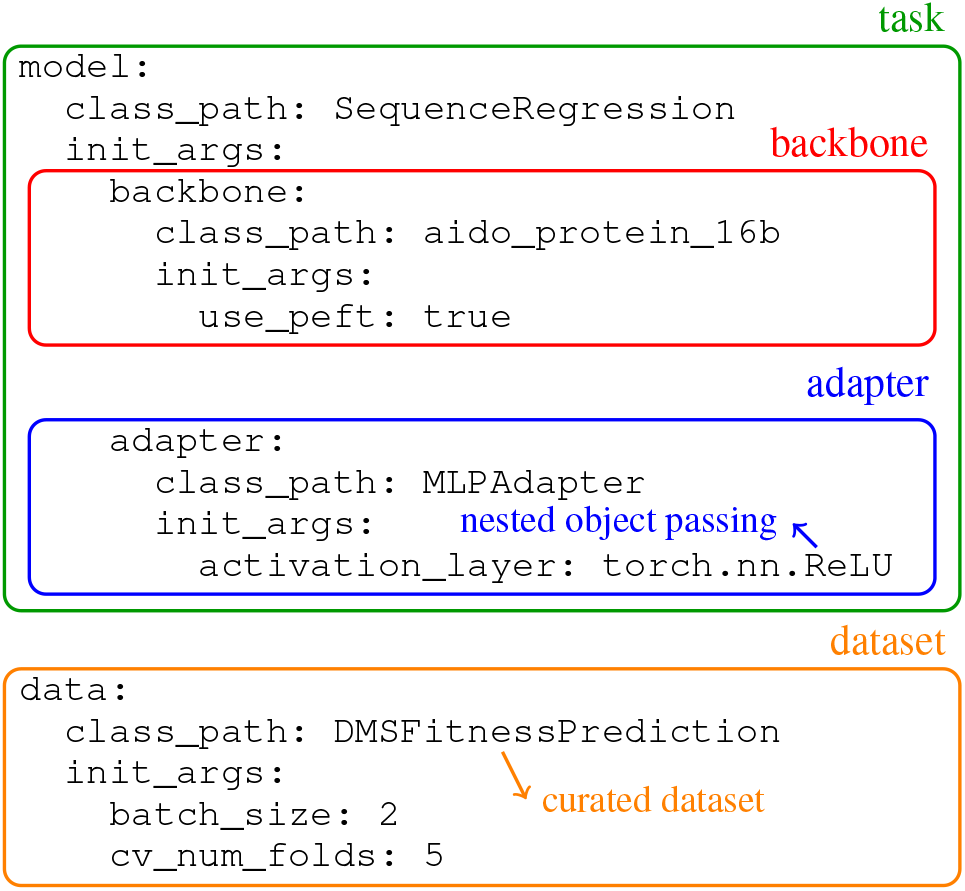
Example configuration with annotations illustrating how each component is configured independently.

~~~
mgen fit/test/predict --config config.yaml
~~~

For every `mgen` call, a full YAML is logged with all user-specified and default arguments, ensuring full reproducibility without direct oversight. Together, this enables quick no-code development and testing. Combined with containerization and Lightning’s deterministic run option, this ensures one-command byte-for-byte reproducibility across machines and time.

**Backbones** wrap pretrained FMs and all their associated design choices, such as tokenization, weights, and runtime options in an interface which hides unnecessary complexity while exposing critical features for automatic configuration with tasks and adapters. This interface also enables AIDO.ModelGenerator to provide generic small-scale backbones, which reduce computational burdens for developing tasks, adapters, and datasets while still guaranteeing correctness with large-scale models and experiments. Because every backbone is an ordinary `nn.Module`, users can load weights from any Hugging Face repository or local checkpoint with a one-line configuration override. AIDO.ModelGenerator also provides a uniform interface for parameter-efficient finetuning (PEFT) on Hugging Face-based backbones (Mangrulkar et al., 2022). When using PEFT, efficient checkpointing can also be enabled, wasting no disk space by only saving unfrozen weights.

**Tasks** define data and model usage. They provide a simple interface for swapping backbones, adapters, and data without any code changes, enabling rapid and reproducible experimentation and leaderboard creation. Tasks are comparable to Hugging Face’s `peft.TaskType`, but cover a wider range of use-cases and data types for information extraction, domain adaptation, supervised prediction, generative modeling, and zero-shot applications.

**Adapters** are model heads that are automatically configured to connect backbones to tasks for prediction or loss evaluation. Adapters are designed for interoperatbility across backbones and tasks (e.g. supervised prediction, conditional generation), avoiding redundant implementations as seen with Hugging Face heads and reducing the risk of bugs. In benchmarking studies, this also decouples the effect of model pretraining from finetuning architectures, enabling fair evaluation of design choices in both stages with minimal overhead. Existing adapters cover popular finetuning architectures, such as transformers, ResNets, cross-attention, MLPs, and linear heads, but are simple to extend and modify as `nn.Module` components.

**Datasets** implement a data interface that extends Lightning’s DataModules to enable automatic loading of remote and local datasets using Hugging Face’s datasets package, as well as automatic train/validation/test splitting and k-fold cross-validation.

**Table 1.** AIDO.ModelGenerator systematically benchmarks modalities’ individual importance and their synergies on the task of predicting tissue-specific RNA isoform expression. Reported results are the maximum from any pre-trained model tested in each category. ^†^Garau-Luis et al. (2024).

| Model | $R^2$ Score ↑ | Spearman ↑ |
| --- | --- | --- |
| Protein | 0.316 | 0.507 |
| RNA | 0.365 | 0.560 |
| DNA | 0.445 | 0.656 |
| RNA + Protein | 0.379 | 0.573 |
| DNA + Protein | 0.465 | 0.674 |
| DNA + RNA | 0.584 | **0.734** |
| DNA + RNA + Protein | **0.586** | **0.734** |
| Isoformer<sup>†</sup> | 0.446 | 0.714 |

### 2.1. Multimodal Fusion

To interrogate cross-modal biology, we provide fusion tasks which compose multiple backbones into a single model. Because each backbone conforms to the same interface, any combination of DNA, RNA, protein, structure, single-cell, or spatial models can be wired together declaratively.

~~~
model:
  class_path: MMSequenceRegression
  init_args:
    backbone: enformer
    backbone1: aido_rna_650m
    backbone2: esm2_15b
    fusion: CrossAttentionFusion
    adapter: MLPAdapter
~~~

## 3. Use Cases

AIDO.ModelGenerator lowers the barrier-to-entry for FM usage while also providing advanced multimodal functionality. For machine learning researchers, we demonstrate a benchmarking study for multimodal fusion of large-scale pretrained FMs, leading to a new SOTA for tissue-specific RNA isoform expression prediction. For biologists, we demonstrate that AIDO.ModelGenerator provides a simple and effective way to use FMs to go beyond classical biological analyses by using it to simulate biological systems for drug discovery. Full experiment results and reproducible configurations are available online ^2 3^.

**Table 2.** AIDO.ModelGenerator substitutes backbones for a given data type in unimodal and multimodal tasks, systematically selecting the best pretrained model for a given task. ^†^AIDO.RNA-CDS 1.6B. ^‡^Garau-Luis et al. (2024).

| DNA Model | RNA Model | $R^2$ Score ↑ | Spearman ↑ |
| --- | --- | --- | --- |
| × | RNA-FM | 0.325 | 0.514 |
| × | RiNALMo | 0.361 | 0.554 |
| × | AIDO.RNA<sup>†</sup> | 0.365 | 0.560 |
| Enformer | RNA-FM | 0.569 | 0.721 |
| Enformer | RiNALMo | **0.585** | 0.733 |
| Enformer | AIDO.RNA<sup>†</sup> | 0.584 | **0.734** |
| Isoformer<sup>‡</sup> |  | 0.446 | 0.714 |

### 3.1. Multimodal Fusion

We apply AIDO.ModelGenerator to systematically benchmark unimodal and multimodal modeling approaches on the tissue-specific isoform expression benchmark proposed by Garau-Luis et al. (2024). Each sample comprises 190kbp DNA, the RNA isoform of interest, and the translated protein (if coding). The task is to regress log_2_(0.0001+TPM) for 170k isoforms across 30 tissues. We apply AIDO.ModelGenerator to answer 3 questions: (1) How important is each modality for this task? (2) Does multimodal modeling improve performance? (3) Which pretrained FM is best for this task?

We apply Enformer (Avsec et al., 2021) and AIDO.DNA (Ellington et al., 2024) for DNA representation, AIDO.RNACDS 1.6B (Zou et al., 2024), RNA-FM (Chen et al., 2022), and RiNALMo (Penić et al., 2024) for RNA representation, and ESM2 (Lin et al., 2023) for protein representation. To assess modality importance, we systematically cycle through all single, double, and triple modality combinations for these models, reporting the best scores for each combination (Fig. 1). AIDO.ModelGenerator identifies that DNA is the single most important modality, followed by RNA and finally protein. Multimodal fusion of DNA+RNA improves performance by over 10% compared to any single modality model, but additional protein data provides no extra benefit.

We then assess the efficacy of RNA models with and without the use of DNA in tandem. We fix Enformer and cycle through AIDO.RNA-CDS 1.6B (Zou et al., 2024), RNA-FM (Chen et al., 2022), and RiNALMo (Penić et al., 2024), achieving a SOTA 0.734 Spearman with Enformer and AIDO.RNA.

### 3.2. Target Identification for Crohn’s Disease

To demonstrate how AIDO.ModelGenerator facilitates plug-and-play functionality within existing bioinformatics pipelines, we apply it to identify therapeutic targets for Crohn’s disease. Our dataset consists of 117,912 scRNA-seq measurements on ileal biopsies from treatment-naive Crohn’s disease (CD) patients and healthy controls (Garrido-Trigo et al., 2023). We evaluated methods in terms of their ability to prioritize the target SOX4, a target of known interest based on genomic evidence for its role in intestinal inflammation (Karczewski et al., 2020).

**Table 3.**
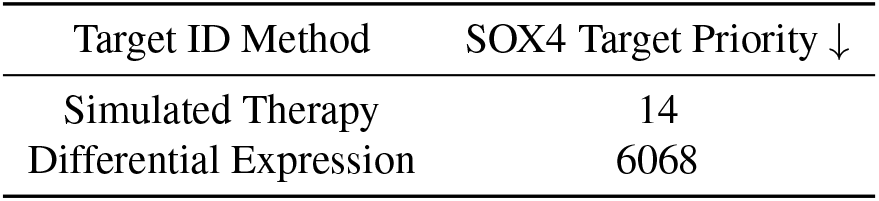
Ranking SOX4, a known target for Crohn’s disease by OpenTargets, according to FM-based and traditional target identification analyses.

We first performed a conventional differential-expression (DE) screen using scanpy (Wolf et al., 2018), which scored SOX4 at 6,068 out of 18,826 total genes. Then, we used AIDO.ModelGenerator to simulate a therapeutic that knocked down each gene and prioritized targets based on the predicted efficacy of this hypothetical therapeutic. We simulated a therapy by setting the expression of each gene in each disease cell to zero, then used AIDO.ModelGenerator to call AIDO.Cell (Ho et al., 2024) to embed these virtually drugged cells as well as the healthy cells. We ranked targets according to the inverse L2 distance between the drugged cells and the healthy cells, favoring knockouts that make diseased cells “look healthy” in embedding space (Fig. 3) Here, SOX4 is prioritized as the 14th most effective target.

## 4. Discussion and Impact

AIDO.ModelGenerator collapses weeks of multimodal FM plumbing into a standardized YAML-driven workflow, providing minimal-overhead interfaces that link an ever-expanding set of 300+ datasets and 30+ backbones to reproducible and engineerable workflows. Ablation and target-identification studies display AIDO.ModelGenerator’s ability to facilitate both machine learning and biology-oriented studies, providing a shared language for cross-disciplinary research teams in BioML. The current release favors sequence-adjacent data types, lightweight fusion heads and adapters, and common parallelism strategies; upcoming work will add deeper integration with imaging, health, and multi-omics data types and more sophisticated parallelisms for multi-FM fusion. By providing a hackable and reproducible framework for development with FMs, AIDO.ModelGenerator seeks to accelerate translation of multimodal FMs from prototype to practice.

## A. Extended Results

**Figure 3.**
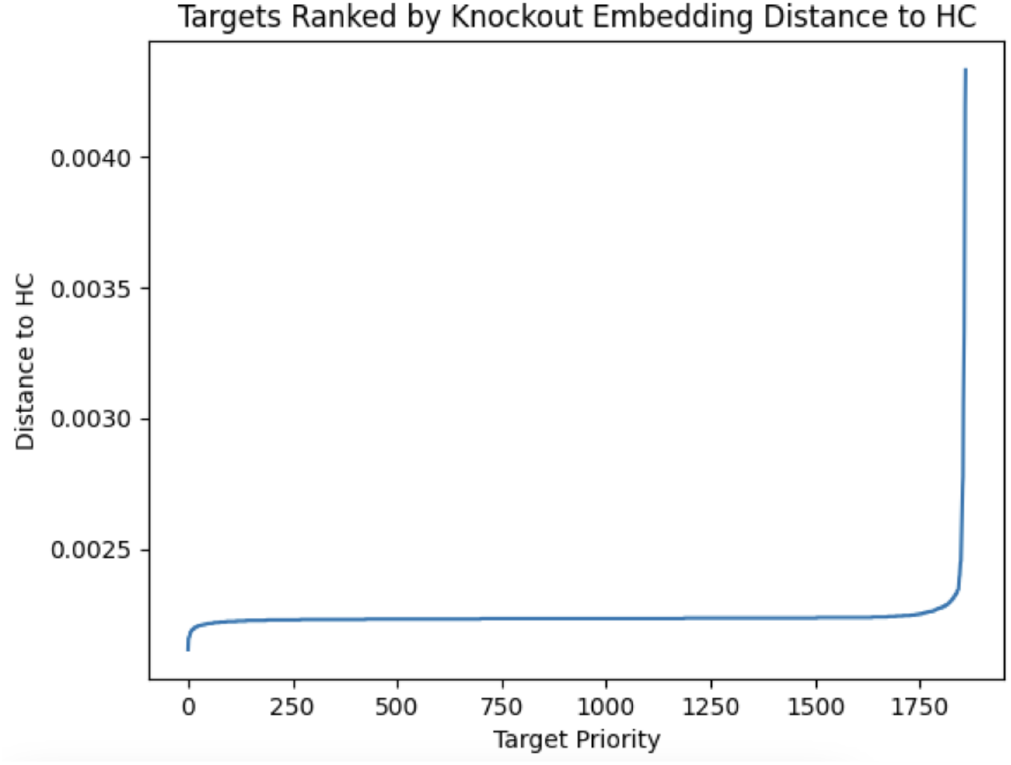
Using AIDO.ModelGenerator to apply AIDO.Cell to target pioritization by simulating therapeutic activity through gene knockouts. Genes are assigned a priority based on how closely they resemble a healthy cell (HC) embedding when they are set to zero, simulating a therapy in a diseased cell.

## Footnotes

1 https://genbio-ai.github.io/ModelGenerator/

2 https://github.com/genbio-ai/ModelGenerator/tree/main/experiments/AIDO.RNA/multimodal_isoform_expression

3 https://github.com/genbio-ai/ModelGenerator/blob/main/experiments/AIDO.Cell/tutorial_target_id.ipynb

